# Intravenous Immunization with Triple Auxotrophs of *Mycobacterium tuberculosis*: A novel vaccine strategy against tuberculosis

**DOI:** 10.1101/2024.05.15.594337

**Authors:** Catherine Vilchèze, Saranathan Rajagopalan, William R. Jacobs

## Abstract

Tuberculosis, caused by *Mycobacterium tuberculosis* (*Mtb*), remains a leading infectious cause of mortality worldwide despite widespread use of the BCG vaccine and the availability of sterilizing pharmacopoeia. Recent research indicates that the intravenous administration of BCG confers sterilizing immunity against *Mtb* pulmonary challenge in non-human primates. However, while BCG is relatively safe, complications such as disseminated BCGosis have been observed in immunocompromised individuals. Double auxotrophic mutants of *Mtb* lacking the ability to synthesize leucine and pantothenate are safe and sterilized in immunocompromised mice and SIV-infected Rhesus macaques. We examined how immunization with a *Mtb* triple auxotrophic strain, mc^2^7902, which cannot synthesize leucine, pantothenate, and arginine, protects immunocompetent mice from a virulent *Mtb* infection. The route of immunization was a crucial factor for protection with mc^2^7902 with intravenous immunization being 100 times more effective in protecting immunocompetent mice from *Mtb* challenge when compared to conventional subcutaneous vaccination with BCG. To further increase the safety of the attenuated auxotroph for vaccine purposes, the type VII secretion system Esx1 responsible for BCG attenuation was deleted in mc^2^7902. When tested by prime-boost immunization of immunocompetent mice followed by aerosol challenge with virulent *Mtb*, mc^2^7902 Δ*esx1* provided similar protection to mc^2^7902. This robust protection against *Mtb* infection conferred by mc^2^7902 and mc^2^7902 Δ*esx1* in a mouse model paves the way for new TB vaccine development using highly attenuated, auxotrophic *Mtb* strains.

## Introduction

Tuberculosis (TB) causes over 1.3 million deaths and over 10 million TB cases per year (1). These infections occur despite the availability of BCG, the only licensed TB vaccine, and antimicrobial therapy that can sterilize *Mycobacterium tuberculosis* (*Mtb*), the causative agent of TB. BCG is a live, attenuated vaccine, derived from *Mycobacterium bovis* (2). BCG lost its virulence through multiple passages that yielded a 9.5 kbp genomic deletion, which partially encodes the type VII secretion system Esx1, a key virulence system in *Mtb*-host interactions (3). While generally safe for healthy individuals, BCG can cause serious life-threatening disease among individuals with compromised immune systems (4-6). The BCG vaccine is currently administered intradermally, but BCG was initially developed as an oral vaccine (2). The route of vaccination triggers different immune responses and is therefore an important factor in immunization studies (7-9). Recent studies showed that intravenous (iv) immunization of non-human primates with BCG confers sterilizing immunity against challenge with virulent *Mtb* (7, 9, 10), although only a modest improvement in protection was observed in the mouse challenge model (11).

We previously demonstrated that *Mtb* auxotrophic strains are safe in animals and provide protection against TB (12-14). We constructed a triple auxotrophic *Mtb* strain for biosynthesis of the vitamin pantothenate and amino acids leucine and arginine [mc^2^7902, H37Rv Δ*leuCD* Δ*panCD* Δ*argB*] (15). mc^2^7902 does not kill immunocompetent or immunocompromised mice and is cleared in both mice within six months (16). Due to its demonstrated safety in mice, we tested whether mc^2^7902 could be a viable TB vaccine candidate. We present data showing significant protection of immunocompetent mice immunized intravenously with mc^2^7902 and mc^2^7902Δ*esx1* when challenged with virulent *Mtb*.

## Results

The protective efficacy of mc^2^7902 was tested through three routes of immunization: subcutaneous (SQ), iv, and orally (gavage). Immunocompetent C57BL/6 mice were primed and boosted three weeks apart with BCG SQ or with mc^2^7902 (SQ, iv, or oral) followed by aerosol challenge with *Mtb* H37Rv three weeks after the boost. We did not observe any protection when mc^2^7902 was given orally (Fig. 1A). SQ immunization with either mc^2^7902 or BCG Danish resulted in similar levels of protection: no more than a log reduction in *Mtb* H37Rv burden in mouse lung, spleen, and liver. In contrast, the H37Rv burden was 100 times lower in the lungs of mice immunized with mc^2^7902 iv compared to SQ three weeks post challenge, with bacteria below the limit of detection in spleen and liver. Eight weeks post challenge, bacterial load had increased in organs of mice with iv mc^2^7902 immunization but was still lower than for mc^2^7902 SQ immunization.

**Figure 1.**
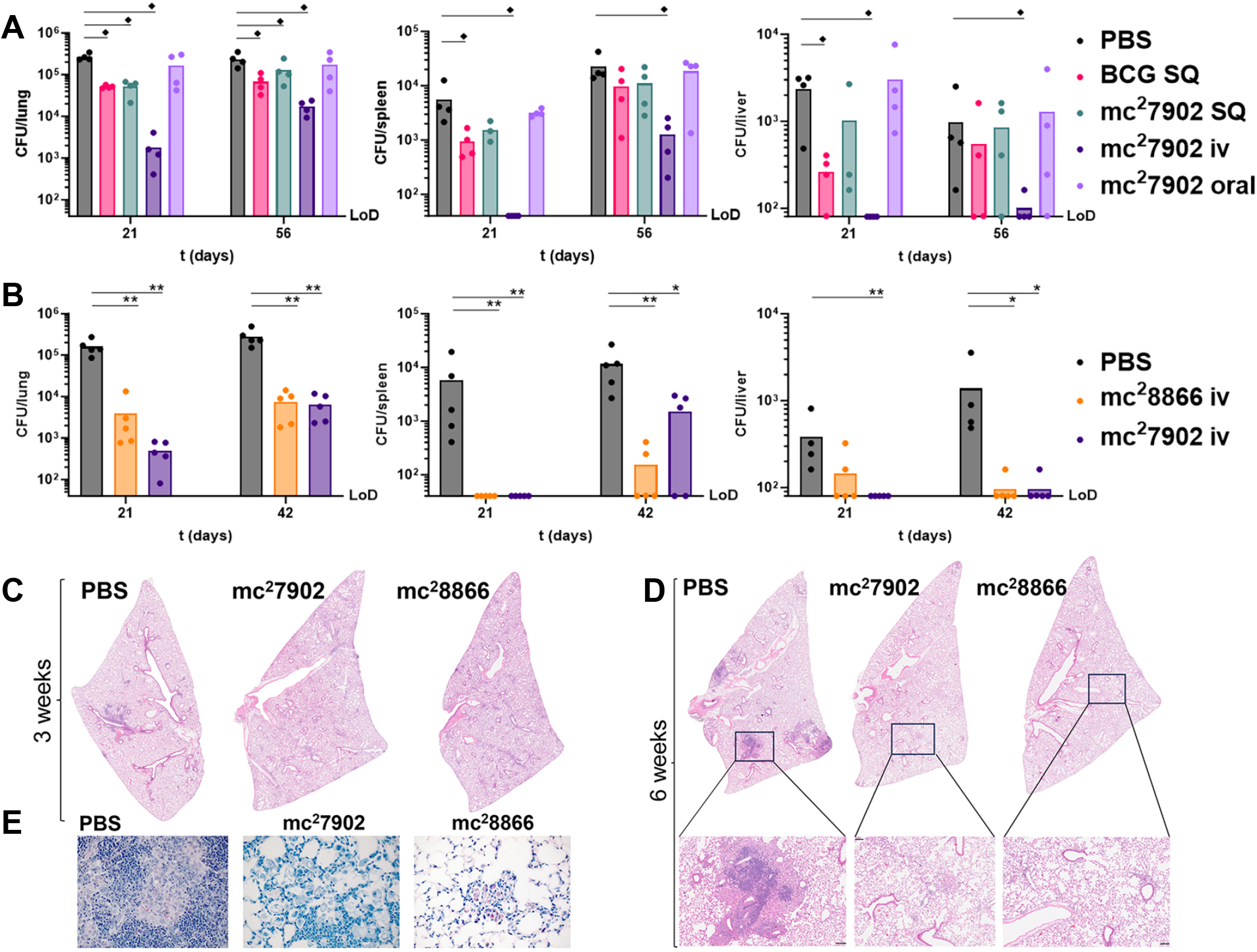
Intravenous immunizations with mc^2^7902 and mc^2^8866 protect C57BL/6 mice from *Mtb* infection. *Mtb* H37Rv bacterial loads in lung (left panel), spleen (middle panel), and liver (right panel) of mice primed and boosted three weeks apart with: (*A*) BCG Danish SQ (5×10^5^ CFUs/injection), mc^2^7902 SQ (5×10^5^ CFUs/injection), mc^2^7902 iv (5×10^6^ CFUs/injection), or mc^2^7902 orally (5×10^6^ CFUs/gavage), and challenged three weeks after the boost with H37Rv (235 CFUs; 4 mice per group and per time point); or (*B*) mc^2^7902 iv (4×10^6^ CFUs/injection) or mc^2^8866 iv (mc^2^7902 Δ*esx1*, 4×10^6^ CFUs/injection) and challenged three weeks after the boost with H37Rv (188 CFUs; 5 mice per group and per time point). Lung, spleen, and liver were homogenized and plated to determine colony-forming-units (CFUs) in each organ. Dots represent CFUs for one mouse, and each bar represents the mean of the data (LoD, limit of detection). Statistically significant data are marked with ^♦^ (p 0.03), * (p 0.016), or ** (p 0.008). (*C, D*) Representative photomicrographs of hematoxylin/eosin-stained lung sections at 3 weeks (*C*) and 6 weeks (*D*) post H37Rv challenge from experiment shown in panel *B* (magnification 20×, bar = 0.1 mm). (*E*) Acid-fast staining of lung sections at 6-week post H37Rv challenge from experiment shown in panel *B* (magnification 40×).

Pursuing the remarkable protection offered by iv immunization with mc^2^7902, we examined whether deleting genes encoding the type VII secretion system Esx-1 would further improve the protective efficacy of mc^2^7902. The partial loss of the Esx1 region attenuated BCG, suggesting that mc^2^7902 Δ*esx1* (mc^2^8866) would be an even safer strain than mc^2^7902. Furthermore, vaccination with mc^2^8866 would not interfere with the interferon-gamma release assay (IGRA), which uses the Esx1 secretion system antigens ESAT-6 and CFP-10 to detect latent TB.

Following the same prime-boost and aerosol challenge protocol of our first experiment, we confirmed the protection by mc^2^7902 iv immunization, observing a 2-log reduction in lung bacterial load three weeks post challenge compared to non-immunized mice (Fig. 1B). Consistent with previous findings, *Mtb* was below the limit of detection in spleens and liver of mc^2^7902 mice immunized intravenously. Protection with mc^2^8866 was not superior to mc^2^7902 three weeks post challenge (Fig. 1B), but organ bacterial burden did not increase as quickly between weeks 3 and 6 post *Mtb* challenge with mc^2^8866 versus mc^2^7902. However, while all lung samples at the 3-week time point showed several small lesions (Fig. 1C), lesions became rare and very small by 6 weeks for mc^2^7902– and mc^2^8866-immunized mice (Fig. 1D). Furthermore, perivascular/peribronchiolar lymphoid infiltrates were mild to moderate in all lung samples at 3 weeks and became mild (for mc^2^7902) and minimal to mild (for mc^2^8866) by 6 weeks (Fig. 1D). Acid-fast bacilli were not found in the lungs of mc^2^7902-immunized mice at 3– and 6-weeks post challenge and rarely in mc^2^8866-immunized mice at 6 weeks (Fig. 1E). Overall, our findings suggest that iv immunization with mc^2^7902 or mc^2^8866 results in improved protection compared to BCG after H37Rv challenge.

## Discussion

Our study demonstrates a remarkable level of vaccine protection that opens new avenues for investigating protective immune responses against TB in mouse models. While sterilizing immunity was observed against *Mtb* among BCG iv-vaccinated primates, *Mtb* elimination in vaccinated mice was not achieved except with ultra-low dose *Mtb* infection of SQ BCG-vaccinated mice (17). While there was only a modest increased in protection in mice vaccinated with BCG intravenously compared to intradermal (11), the route of immunization with mc^2^7902 was central to the protection of *Mtb* infected mice: intravenous immunization with mc^2^7902 offered a 50-100-fold better protection than SQ immunization. A previous study showed that BCG iv immunization in mice induced a “trained immunity” phenotype where iv BCG, but not SQ BCG, yielded transcriptional changes in hematopoietic stem cells resulting in the increased ability of macrophages to protect against *Mtb* infection (18). The immunological underpinnings of our enhanced protection, although not understood, could be caused by a rewiring of the macrophage’s ability to fight *Mtb* infection, as observed during iv BCG immunization (18), or possibly by the high *Mtb* cell numbers reaching the spleen during iv immunization, yielding an enhanced immune response.

Using iv immunization in humans would necessitate extensive safety studies. However, mc^2^7902 has non-revertible and non-suppressible deletions of five genes (*panC, panD, leuC, leuD, argB*) and is safe in immunocompromised mice, providing a strong foundation for further development as a vaccine. mc^2^7902 does not persist in immunocompetent and immunocompromised mice and is eliminated in the lung, liver and spleen of the infected mice after six months (16), suggesting that mc^2^7902 could be a safe vaccine in immune-challenged individuals. Furthermore, the lung inflammation in the infected mice immunized with mc^2^7902 or with mc^2^8866 seems to decrease over time, signaling a reduced disease burden although this was not correlated by the CFUs data. Integrating additional immune-evasion mutations in mc^2^7902 could further augment its immunogenicity, leading to the development of a more effective vaccine for human TB and potentially for *M. bovis* infection in animals (19). Our future studies include using non-human primate models of TB to evaluate the vaccine efficacy of mc^2^7902 and derivative strains and to identify specific components of the immune response that contribute to effective protection by TB vaccines.

## Methods

### Bacterial strains, media, and materials

The mycobacterial strains used in this study were obtained from laboratory stocks. The strains were grown shaking at 37°C in Middlebrook 7H9 supplemented with 10% (v/v) OADC enrichment (Difco, Sparks, MD), 0.2 % (v/v) glycerol, and 0.05% (v/v) tyloxapol. For mc^2^7902 and mc^2^7902 Δ*esx1*, 50 mg/l leucine, 24 mg/l pantothenate and 1 mM arginine were added to the media. Chemicals and biologicals were obtained from Sigma (St Louis, MO) or Thermo Fisher Scientific (Waltham, MA). For *Mtb* H37Rv CFU determination, plating was done on Middlebrook 7H10 supplemented with 10% (v/v) OADC enrichment and 0.2 % (v/v) glycerol (7H10 plate); and plates were incubated at 37°C for 4 to 8 weeks.

### mc^2^7902 Δ*esx1* construction

*esx1* was deleted from mc^2^7902 using specialized transduction (20). Briefly, the left and right flanks of *esx1* were PCR-amplified using the following primers: Esx-1 LL (TTTTTTTTCCATAAATTGGAACCATCAGCGGACCTGTC GGAC), Esx-1 LR (TTTTTTTTCCATTTCTTGGCGCTCATTAGAATAAGTCGGGATGTCTCACTGAGGTCTCTTTATGTGTTTCCTTACGCTCGCCGTTC), Esx-1 RL (TTTTTTTTCCATAGATTGGATCAGGTCTAGGTGAGCATCCGAGTGTCTGGTCTCGTAGTCCCAGAACACTCCATTCGTTGAGATTC) and Esx-1 RR (TTTTTTTTCCATCTTTTGGCGGTGGAGGGGCAGACCAAC). The PCR fragments were digested with Pflm1 cloned into Pflm1-cut pYUB1471. The resulting plasmid was cut with pac1, ligated to pac1-cut phAE159, and packaged in vitro (GigapackII, Stratagene, La Jolla, CA). The resulting phasmid was electroporated into mc^2^155. High titer phage lysate was prepared and used to transduce mc^2^7902. Transductants were plated on 7H10 plates containing 75 mg/l hygromycin, 50 mg/l leucine, 24 mg/l pantothenate and 1 mM arginine and checked by 3-primer PCR and whole genomic sequencing. The hygromycin-sacB selection cassette was excised using the phage phAE280 and sucrose selection (20).

### Immunization and challenge experiments

C57Bl/6 female and male mice (6-8 weeks old) were obtained from Jackson Laboratory (Bar Harbor, ME). The animal protocol AUP00001368 titled “Evaluation of the safety and efficacy of attenuated Mycobacterial vaccine vectors” was approved by the Einstein Animal Institute, which is accredited by the “American Association for the Use of Laboratory Animals” (DHEW Publication No. (NIH) 78-23, Revised 1978), and accepts as mandatory the NIH “Principles for the Use of Animals”.

The strains used for the immunization were grown to exponential phase (OD_600nm_ ≈ 0.8 – 1), as described above, pelleted by centrifugation, washed twice with DPBS-tyloxapol (0.05% (v/v)), and sonicated twice for 10 sec. OD_600nm_ was measured and the strains were diluted in Dulbecco’s phosphate-buffered saline containing 0.05% tyloxapol (DPBS-T). The route of immunization was either SQ (target 1×10^7^ CFU/ml, 0.1 ml injection/mouse), tail vein injection (target 5×10^7^ CFU/ml, 0.2 ml injection/mouse) or oral (target 5×10^7^ CFU/ml, 0.2 ml gavage/mouse). For the oral immunization, mice received mc^2^7902 by gavage three times, ten days apart. The non-immunized control group received DPBS-T (0.1 ml) SQ. The mice were boosted three weeks later with similar doses. Three weeks after the boost, the mice were challenged with *Mtb* H37Rv. H37Rv was prepared as described above for the immunization strains and diluted to 1×10^7^ CFU/ml (after OD_600nm_ measurement and based on OD_600nm_ = 1 = 3×10^8^ CFU/ml) prior to aerosol infection. 24 hours after aerosol challenge, five mice from the DPBS-T control group were euthanized, their lungs homogenized and plated undiluted for initial H37Rv infection dose determination. At the indicated times, five mice per group were euthanized, and their spleen, liver and lungs were removed. The organs were homogenized in DPBS, serially diluted in DPBS-T and plated (see above) to determine CFU/organ.

For pathology, lungs were fixed in 10% buffered formalin phosphate. Following paraffin embedment, lung tissues were sectioned and stained with hematoxylin and eosin. Tissues were analyzed by a pathologist at the Histopathology and Comparative Pathology facility at the Albert Einstein College of Medicine.

### Statistics

Differences between groups were analyzed by an unpaired, nonparametric Mann-Whitney test using GraphPad Prism 10.1.0 (San Diego, CA).

## Acknowledgments

W.R.J. acknowledges support from the National Institutes of Health grant AI026170. We thank Bing Chen, Mei Chen, and John Kim for technical assistance with the mouse experiments and Riti Sharan for helpful discussion.

